# Adult consequences of repeated nicotine and Δ^9^-tetrahydrocannabinol (THC) vapor inhalation in adolescent rats

**DOI:** 10.1101/2023.09.08.556932

**Authors:** Arnold Gutierrez, Kevin M. Creehan, Yanabel Grant, Michael A. Taffe

## Abstract

The use of Electronic Drug Delivery Systems (EDDS, “e-cigarettes”) to ingest nicotine and Δ^9^-tetrahydrocannabinol (THC) has surged in adolescent populations in the United States, as five times as many high-school seniors vape nicotine daily as use tobacco. At the same time 19.5% of seniors use cannabis at least monthly, with 12% using EDDS to deliver it. This study was conducted to examine the impact of repeated adolescent vapor inhalation of nicotine and THC in rats.

Female Sprague-Dawley rats were exposed to 30-minute sessions of vapor inhalation, twice daily, from Post-Natal Day (PND) 31 to PND 40. Conditions included vapor from the propylene glycol (PG) vehicle, Nicotine (60 mg/mL in the PG), THC (100 mg/mL in the PG) or the combination of Nicotine (60 mg/mL) and THC (100 mg/mL). Rats were assessed on wheel activity, heroin anti-nociception and nicotine and heroin vapor volitional exposure during adulthood.

Nicotine exposed rats exhibited few differences as adults, but were less sensitive to anti-nociceptive effects of heroin (1 mg/kg, s.c.). THC- and THC+Nicotine-exposed rats were less spontaneously active, and obtained fewer nicotine vapor deliveries as adults. In contrast, THC exposed rats obtained volitional heroin vapor at rates indistinguishable from the non-THC-exposed groups. Repeated THC exposure also caused tolerance to temperature-disrupting effects of THC (5 mg/kg, i.p.).

These studies further confirm that the effects of repeated vapor exposure to THC in adolescence last into early to middle adulthood, including decreased volitional consumption of nicotine. Effects of repeated nicotine in adolescence were comparatively minor.

## 1. Introduction

Popularity and widespread availability of Electronic Nicotine Delivery Systems (ENDS), most commonly referred to as e-cigarettes, is evidenced by survey data in the United States showing that 20-26% of high-school seniors vaped nicotine in the past month 2019-2021 (Miech et al., 2022). Daily nicotine vaping declined from 11.6% in 2019 to 5% in 2020-2021, but rates of daily cigarette smoking were 2-3% across 2019-2021 in these populations (ibid). There is an obvious concern that reductions in the harm associated with cigarette smoking will be at least partially replaced by similar and/or novel harms associated with vaping for nicotine content. Concerningly, pre-teen rates of nicotine use went up during the initial months of COVID-19 related stay-at-home orders, while alcohol use declined (Pelham et al., 2021). A recent review points out the need for scientific research on the health impact of ENDS based exposure to nicotine and furthermore identifies several domains in which adolescents have previously been found to be at increased risk (Fowler et al., 2018). In parallel with the use of cigarettes and ENDS to ingest nicotine, significant numbers of human adolescents exhibit high rates of cannabis use. Approximately 6% of high-school seniors report using cannabis daily/near daily, and 20-23% endorsed past-month use, from 1997-2021; with 12% reporting that they vaped cannabis in the past month in 2020-2021 (Johnston et al., 2021; Miech et al., 2022). Some individuals use ENDS devices for ingesting opioids (Blundell, Dargan and Wood, 2018), however there is a lack of specific study on these practices (Morris, Pebley and Little, 2023). Thus, these systems are more accurately termed Electronic *Drug* Delivery Systems (EDDS).

The goal of this study was to determine any lasting impact of repeated vapor inhalation of nicotine and THC in adolescent rats on the effects of nicotine and heroin in adulthood. For this, we used an EDDS-based system that has been previously shown effective in delivering nicotine to adult rats (Javadi-Paydar et al., 2019b; Lallai et al., 2021; Montanari et al., 2020), adolescent rats (Gutierrez et al., 2022) and to rat pups in utero (Breit et al., 2022; Hussain, Breit and Thomas, 2022). EDDS-based systems have shown efficacy for nicotine self-administration in rats (Lallai et al., 2021; Smith et al., 2020) and mice (Cooper, Akers and Henderson, 2021; Henderson and Cooper, 2021), and have been used to show that repeated daily exposure to nicotine vapor leads to significant withdrawal following discontinuation in adult rats (Montanari et al., 2020). We have shown that repeated exposure to the major psychoactive constituent of cannabis, Δ^9^-tetrahydrocannabinol (THC), via vapor inhalation during adolescence in rats produces lasting consequences in adulthood. This included alterations in hypothermic and anti-nociceptive effects of THC in both sexes, increases food consumption in male rats and increased self-administration of fentanyl in female rats (Nguyen et al., 2020b). Similarly, twice daily vapor exposure to nicotine from PND 31-40 led to lasting alterations in the self-administration of nicotine by vapor inhalation in female, but not male, rats (Gutierrez et al., 2022). Thus, the overall approach of twice-daily adolescent vapor inhalation in brief sessions (30 minutes) using female rats was selected for this investigation.

This study exposed groups of adolescent female rats to twice daily inhalation of vapor from the propylene glycol (PG) vehicle, including nicotine (60 mg/mL in the PG), THC (100 mg/mL in the PG) or the combination of nicotine (60 mg/mL) and THC (100 mg/mL) for 10 days to determine lasting effects in adulthood. Rats were evaluated in adulthood for thermoregulatory and anti-nociceptive responses to THC and heroin, spontaneous locomotor activity on a wheel, the effect of nicotine on wheel activity, as well as on volitional nicotine and heroin exposure.

## 2. Methods

### 2.1 Subjects

Female Sprague-Dawley (Envigo/Harlan) rats (N=32) were used for this study. The vivarium was kept on a 12:12 hour reversed light-dark cycle, and behavior studies were conducted during the vivarium dark period. Food and water were provided ad libitum in the home cage. Animal body weights were recorded weekly, beginning at 6 weeks of age (PND 36) and continuing through the end of the study. Experimental procedures were conducted in accordance with protocols approved by the Institutional Animal Care and Use Committee of the University of California, San Diego and consistent with recommendations in the NIH Guide (Garber et al., 2011).

### 2.2 Drugs

Nicotine bitartrate (Sigma Pharmaceuticals LLC; North Liberty, IA) or heroin HCl (NIDA Drug Supply) were dissolved in propylene glycol (PG) for vapor inhalation studies and dissolved in physiological saline for subcutaneous injection studies. Concentrations of 30 and 60 mg/mL in the Propylene Glycol (PG) vehicle were used for nicotine and 50 mg/mL for heroin. THC (100 mg/mL) was suspended in PG at a concentration of 100 mg/mL for vapor inhalation studies and suspended in a 1:1:18 (ethanol:cremophor:saline) vehicle for intraperitoneal injection studies. PG was used as the vapor vehicle for consistency and comparability with our prior reports on the impact of nicotine, THC and heroin vapor inhalation (Gutierrez, Creehan and Taffe, 2021; Javadi-Paydar et al., 2019a; Javadi-Paydar et al., 2019b; Taffe et al., 2021a).

### 2.3 Apparatus

#### 2.3.1 Vapor Inhalation

An e-cigarette based vapor inhalation system (La Jolla Alcohol Research, Inc) which has previously been shown to deliver active doses of THC, cannabidiol, heroin, oxycodone, methamphetamine and nicotine (Gutierrez, Creehan and Taffe, 2021; Javadi-Paydar et al., 2019a; Javadi-Paydar et al., 2019b; Nguyen et al., 2016a; Nguyen et al., 2019) was used for these studies. Vapor was delivered into sealed vapor exposure chambers (152 mm W X 178 mm H X 330 mm L; La Jolla Alcohol Research, Inc, La Jolla, CA, USA) through the use of e-vape controllers (Model SSV-3 or SVS-200; 58 watts, 0.24-0.26 ohms, 3.95-4.3 volts, ∼214 °F; La Jolla Alcohol Research, Inc, La Jolla, CA, USA) to trigger SMOK Baby Beast Brother TFV8 sub-ohm tanks. Tanks were equipped with V8 X-Baby M2 0.25 ohm coils. MedPC IV software was used to schedule and trigger vapor delivery (Med Associates, St. Albans, VT USA). The apparatus and settings were the same for all drug conditions. The chamber air was vacuum-controlled by a chamber exhaust valve (i.e., a “pull” system) to flow room ambient air through an intake valve at ∼1 L per minute. This also functioned to ensure that vapor entered the chamber on each device triggering event. The vapor stream was integrated with the ambient air stream once triggered. Airflow was initiated 30 seconds prior to, and discontinued 10 seconds after the initiation of, each puff. Each vapor puff delivery totaled six seconds in duration, and there were a total of six puffs delivered (i.e., at 0, 5, 10, 15, 20 and 25 minutes).

#### 2.3.2 Activity Wheels

Experimental sessions were conducted in white illuminated procedure rooms with activity wheels that attached to a typical housing chamber with a door cut to provide access to the wheel (Med Associates; Model ENV-046), using approaches previously described (Gilpin et al., 2011; Miller et al., 2013; Taffe et al., 2021b). Rats were given access to the wheel in acute 30-minute sessions during which wheel rotation (quarter-rotation resolution) activity was recorded at 10-minute intervals. One habituation session was conducted for each animal prior to initiating the following experimental sessions.

### 2.4 Experiments

#### 2.4.1 Experiment 1: Effect of repeated adolescent nicotine and THC inhalation on nociception and body temperature

Groups (N=8 per group) of female Sprague-Dawley rats arrived in the laboratory at PND25 and were exposed to 30-minute vapor sessions in white light (during the vivarium dark cycle) twice per day (6h apart; 800/1400 hours) for 10 consecutive days from PND 36-45. Vapor conditions included PG, Nicotine (60 mg/mL), THC (100 mg/mL) or the combination of Nicotine (60 mg/mL) and THC (100 mg/mL) tested in separate groups. The higher concentration of nicotine compared with a prior experiment (Gutierrez et al., 2022) was selected to match the concentration used in JUUL devices and to provide some dose-effect information relative to that prior study. On PND 71 the groups were injected with THC (0.725 mg/kg, i.p.) and assessed on tail withdrawal and rectal temperature before and 30 minutes after injection. On PND 80-81, groups were injected with THC (5.0 mg/kg, i.p.) and assessed on tail withdrawal and rectal temperature before, and 30, 60 and 120 minutes after injection.

#### 2.4.2 Experiment 2: Effect of repeated adolescent nicotine and THC inhalation on wheel activity

From PND 114 onward the groups of female rats from Experiment 1 were assessed for wheel activity in 30-minute sessions. Animals first received a habituation session on one day and then a baseline session three days later, with no prior treatment, to assess any lasting group differences associated with the adolescent vapor treatment conditions. Thereafter the effects of 30-minute inhalation of PG or Nicotine (30 mg/mL; in the dark) were assessed with the order counterbalanced within the groups and a 6-day interval between assessments. Wheel activity without any drug exposure was re-determined PND 150-151.

#### 2.4.3 Experiment 3: Effect of repeated adolescent nicotine and THC inhalation on volitional nicotine vapor consumption

The rats were assessed for volitional exposure to nicotine (30 mg/mL) vapor in 30-minute sessions from PND 155-156 onward. Four chambers with nose-poke manipulanda and two with lever manipulanda were used with the assignment of lever boxes counterbalanced across groups. A response on the drug-associated manipulandum (Fixed Ratio 1 response requirement) resulted in illumination of the cue light and delivery of a 1 second puff of vapor. This was followed by a 20 second timeout during which the cue light remained illuminated and hole/lever responses were recorded, but led to no consequences. Sessions were scheduled no more frequently than every other day since prior work has indicated that sequential days of intravenous nicotine self-administration can produce declining trends (O’Dell and Koob, 2007). Cohorts consisting of half of each of the treatment groups were run on alternating days. A facilities flooding emergency occurred around the tenth (second cohort) and eleventh (first cohort) acquisition sessions and scheduled remediation and repairs disrupted access to testing rooms thereafter. As there may have been effects of the developing water leak in the building, and staff response to it, analysis of the Acquisition phase was limited to the initial 9 sessions. Animals were idled for three weeks and then re-started under the FR1, termed Session 10 here for convenience. After Session 12, the schedule of reinforcement was increased to FR5 for three sessions and then restored to FR1 for three additional sessions. Menthol has been reported to enhance mouse vapor self-administration of nicotine, thus Nicotine 30 mg/mL with 5% menthol was assessed for 3 sessions, followed by Nicotine 60 mg/mL with 5% menthol for an additional 8 sessions. Animals were switched to different operant boxes during the last six of these eight sessions, to determine if individual differences were confounded with the specific operant box.

#### 2.4.4 Experiment 4: Effect of heroin injection on nociception and body temperature

The impact of heroin (0.0, 0.56, 1.0, 1.56 mg/kg, s.c.) on tail withdrawal and rectal temperature was assessed from 42-43 weeks of age (∼PND 296-306) in a counterbalanced order. The approach was as described for Experiment 1.

#### 2.4.5 Experiment 5: Effect of repeated adolescent nicotine and THC inhalation on volitional heroin vapor exposure

Access to volitional vapor was restarted after Experiment 4, at approximately 46-49 weeks of age (∼PND 324-345). Animals were assessed in 30-minute sessions with the opportunity to obtain puffs of heroin (50 mg/mL) vapor under a FR1 schedule of reinforcement for four sessions. These sessions were compared with the final prior nicotine (60 mg/mL + menthol) sessions in the analysis. Tail withdrawal was evaluated before and after the first heroin session.

### 2.6 Data Analysis

Data were analyzed by Analysis of Variance, save that mixed-effect models were used in any cases of missing values, including when percent of responses on the drug-associated manipulandum were undefined due to no drug-associated responses being emitted. Within-subjects factor of Time (after vapor initiation / injection or wheel access), acute treatment condition or self-administration Session, and a between-subjects factor for adolescent treatment group were included as relevant. Due to a main effect of receiving THC during adolescence in the study of rectal temperature after THC administration, subsequent analysis of data for these groups included, *a priori*, an initial four-group assessment using a three factor (Time/Session/acute treatment, Presence / Absence of THC, Presence / Absence of Nicotine) analysis, followed by examining the data in a two factor analysis collapsed across the two groups that received/did not receive THC and then a two factor analysis collapsed across the two groups that received/did not receive nicotine (see **Table 1**), with follow up post-hoc exploration of significant main effects or interactions. In all analyses, a criterion of P<0.05 was used to infer that a significant difference existed. Any significant main effects were followed with post-hoc analysis using Tukey (multi-level factors), Dunnett (comparison with a control condition) or Sidak (two-level factors) correction. All statistical analyses used Prism for Windows (v. 9.5.1-10.0.0; GraphPad Software, Inc, San Diego CA).

**Table 1:**
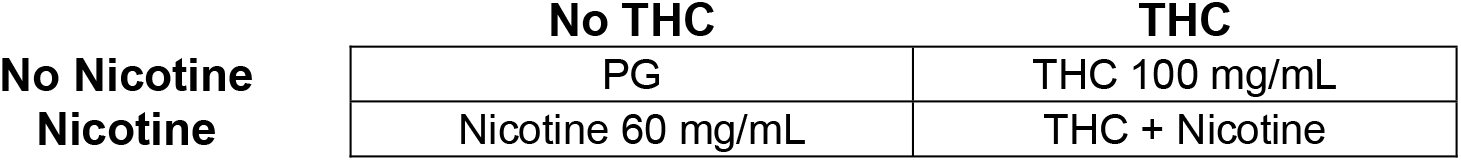
Factorial design for the primary analysis.

## 3. Results

### 3.1 Repeated THC and Nicotine Vapor Exposure

There were no group differences in body weight associated with the adolescent inhalation treatment across the study interval. N=30 completed all scheduled studies and were euthanized between PND 437-444 (∼62-63 weeks of age). One rat (PG group) was observed to develop a clinically concerning solid mass and was euthanized PND 319 and one rat (PG group) was found dead of unknown causes PND 398.

### 3.2 Experiment 1: Effect of repeated adolescent nicotine and THC inhalation on nociception and body temperature

Injection of 0.725 mg/kg THC, i.p., produced threshold effects (not shown); there was a significant effect of Time (before/after THC) for both rectal temperature [F (1, 28) = 24.46; P<0.0001] and tail withdrawal latency [F (1, 28) = 36.30; P<0.0001], but no significant group differences were confirmed in the initial study. The injection of 5 mg/kg THC, i.p., significantly reduced rectal temperature and increased tail withdrawal latencies as is shown in **Figure 1**. The three factor ANOVA confirmed a significant effect of Time [F (1.825, 51.10) = 38.60; P<0.0001] and the interaction of Time with adolescent treatment group [F (3, 84) = 3.37; P<0.05] on rectal temperature. The follow up two factor ANOVA collapsed across the nicotine exposure factor likewise confirmed a significant effect of Time [F (3, 90) = 40.45; P<0.0001] and the interaction of Time with THC/no-THC group [F (3, 90) = 3.54; P<0.05] on rectal temperature. The Tukey post-hoc test further confirmed that temperature was significantly lower than the pre-injection temperature 30, 60 and 120 minutes after injection, within each of the adolescent THC/no-THC groups. The post-hoc test did not, however, confirm any significant differences between the groups at any of the time points.

**Figure 1:**
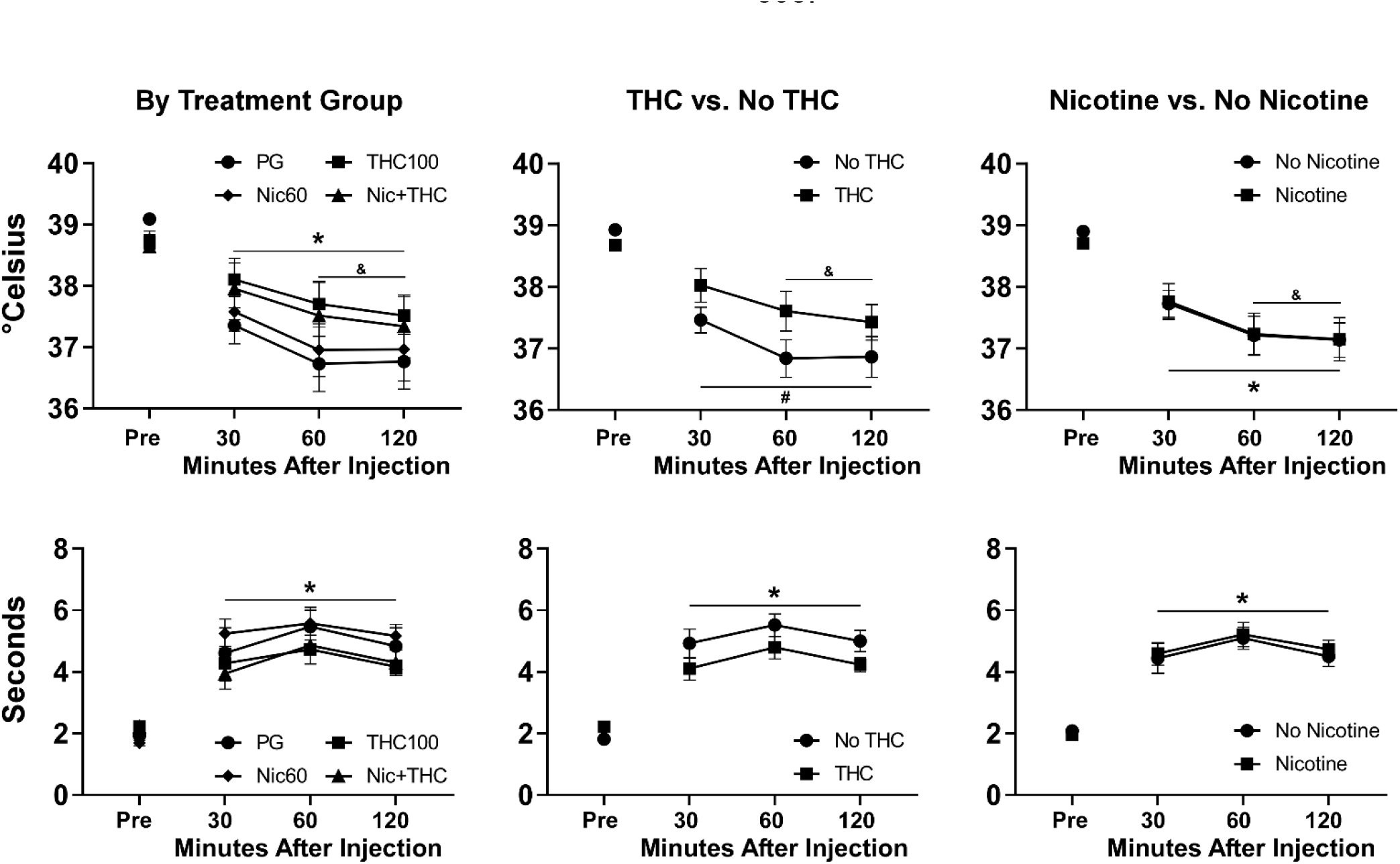
Mean (±SEM) rectal temperature and tail-withdrawal latencies following injection with THC (5.0 mg/kg, i.p.) are depicted for all four treatment groups, for groups which did or did not receive THC and for groups which did or did not receive nicotine. A significant difference from the Pre-injection value, collapsed across group, is indicated with *. A significant difference from the Pre-injection value, within each group, is indicated with #. A significant difference from the 30 minute value, collapsed across group, is indicated with &.

The three-way analysis of the tail withdrawal latencies confirmed a significant effect of Time [F (2.495, 69.87) = 52.71; P<0.0001], but not of adolescent treatment group by itself or in interaction with the Time factor. The post-hoc analysis confirmed that withdrawal latency was slower than the baseline value at every post-injection timepoint. The follow up two-factor analyses using the Nicotine/No-Nicotine groupings did not confirm any significant effects of group on temperature or tail-withdrawal latency.

### 3.3 Experiment 2: Effect of repeated adolescent nicotine and THC inhalation on wheel activity

#### 3.3.1 Baseline test for wheel activity, PND 114

The Baseline (i.e., without any acute drug exposure) wheel activity assessed in early adulthood (PND 114) differed between the two groups that received adolescent vapor exposure to THC (alone or in combination with nicotine) and the two that did not (PG and nicotine only), as depicted in **Figure 2**. The initial three-factor analysis confirmed that there was an effect of Time [F (1.719, 48.14) = 11.51; P<0.0005], and of the presence/absence of THC [F (1, 28) = 4.36; P<0.05], but not of the presence/absence of nicotine, in the adolescent vapor on baseline wheel activity. Post-hoc exploration of the follow-up analysis of the rats grouped by THC/No THC [Time: F (1.689, 50.67) = 11.77; P=0.0001; Group: F (1, 30) = 4.46; P<0.05; Interaction: n.s.] or by Nicotine/No Nicotine [Time: F (1.737, 52.10) = 11.92; P=0.0001; Group: n.s.; Interaction: n.s.] further confirmed significantly less wheel activity in the first 10 minutes for the THC-exposed groups.

**Figure 2:**
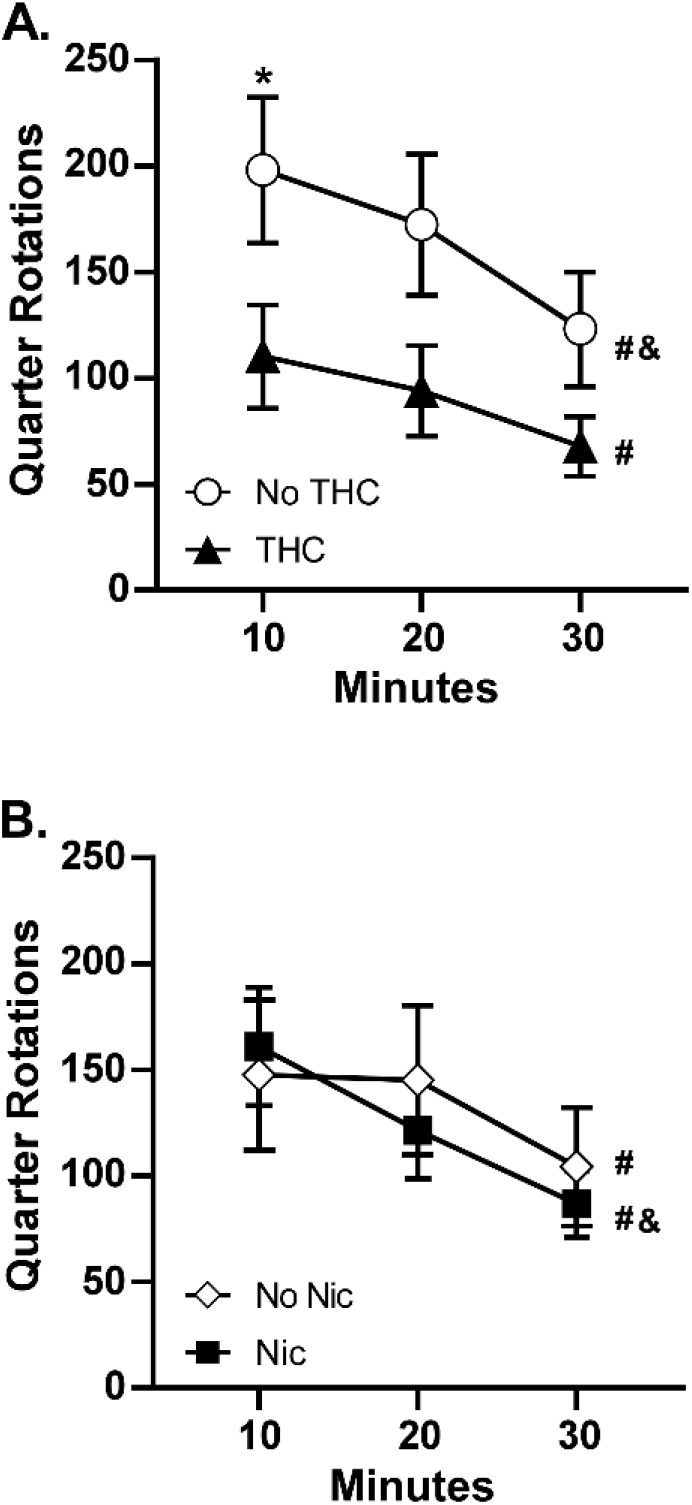
Mean (N=16; ±SEM) quarter rotations of the wheel for the groups that received THC or no THC (A) or that received Nicotine or no Nicotine (B). A significant difference between groups is indicated with *, a difference from the 10 minute bin with # and a difference from the 20 minute bin with &.

#### 3.3.2 Impact of nicotine on wheel activity

Nicotine vapor inhalation before the session significantly suppressed wheel activity (**Figure 3**), as was confirmed by a significant effect of acute treatment condition [F (1.000, 30.00) = 51.99; P<0.0001] in the three factor analysis of the THC vs No-THC groups wheel activity by 10-minute time bin. There were also significant effects of THC/No-THC group [F (1, 30) = 6.44; P<0.05], of Time [F (1.797, 53.90) = 59.60; P<0.0001], as well as of the interactions of Time with THC/No-THC group [F (2, 60) = 4.21; P<0.05] and with the acute treatment condition [F (1.303, 39.10) = 27.73; P<0.0001]. The three-factor analysis of the Nicotine vs No-Nicotine groups confirmed significant effects of Time [F (1.816, 54.49) = 52.30; P<0.0001], of pre-treatment condition [F (1.000, 30.00) = 46.58; P<0.0001] and the interaction of Time with pre-treatment [F (1.000, 30.00) = 46.58; P<0.0001], but there were no significant effects of adolescent treatment group.

**Figure 3:**
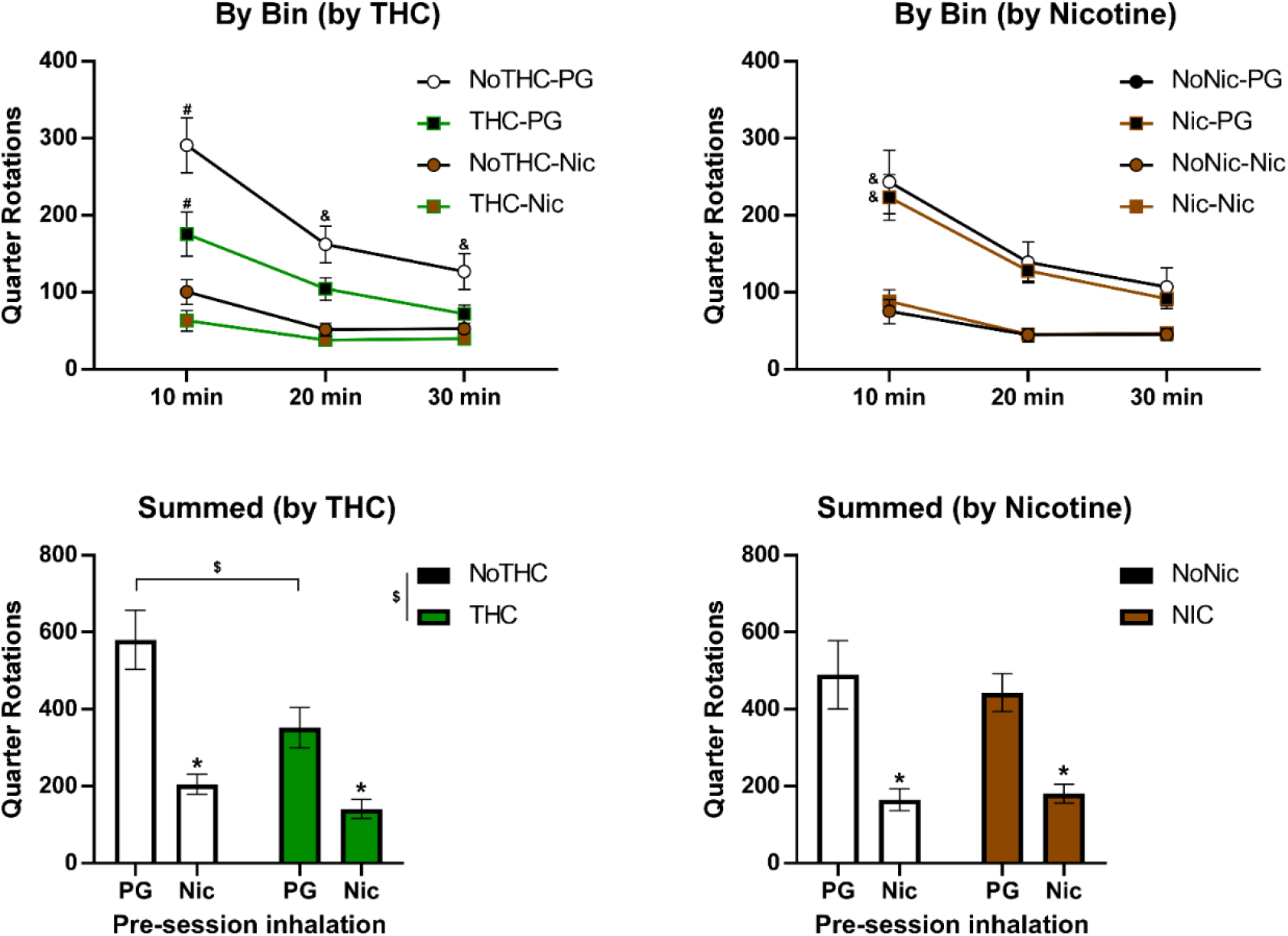
Mean (±SEM) wheel activity following inhalation of nicotine (30 mg/mL) vapor. A significant difference from all other groups/conditions is indicated by # and a difference from each nicotine vapor condition by &. A significant difference between adolescent treatment groups is indicated with $ and significant difference associated with pre-session inhalation condition with *.

Analysis of total session wheel activity with a three factor ANOVA confirmed a significant effect of acute treatment Condition [F (1, 28) = 53.11; P<0.0001], and of the adolescent THC [F (1, 28) = 6.32; P<0.05], however there was no main effect of adolescent Nicotine, nor any interactions with Nicotine confirmed. The follow-up two factor ANOVA of the THC/No-THC grouping again confirmed main effects of Adolescent Treatment [F (1, 30) = 6.44; P<0.05] and of acute treatment condition [F (1, 30) = 51.99; P<0.0001] but there was no significant interaction [F (1, 30) = 4.06; P=0.0529] of factors. The post-hoc test confirmed significant suppressive effects of acute nicotine within each of the THC/No-THC adolescent treatment groupings and a significant difference between adolescent treatment groupings for PG treatment only. The follow-up two factor ANOVA of the Nicotine/No-Nicotine grouping again confirmed main effects of Pre-treatment Condition [F (1, 30) = 51.99; P<0.0001] but there were no significant effects of Nicotine/No-Nicotine adolescent treatment grouping or of the interaction of factors. The post-hoc test again confirmed significant suppressive effects of acute nicotine within each of the Nicotine/No-Nicotine adolescent treatment groupings.

### 3.3.3 Baseline test for wheel activity, PND 150-151

When assessed on PND 150-151 the animals exposed to THC (alone or in combination with nicotine) still emitted less wheel activity than those without THC exposure (PG and Nic alone), as depicted in **Figure 4**. In a three factor ANOVA there were significant effects of Time [F (11, 308) = 70.97; P<0.0001], of THC condition [F (1, 28) = 5.09; P<0.05] and of the interaction of Time with THC condition [F (11, 308) = 1.97; P<0.05]. The post-hoc exploration of the THC/NoTHC groupings further confirmed a significant difference between the groups for the 5 and 10 minute bins. No significant effects of nicotine condition alone or in interaction were confirmed.

**Figure 4:**
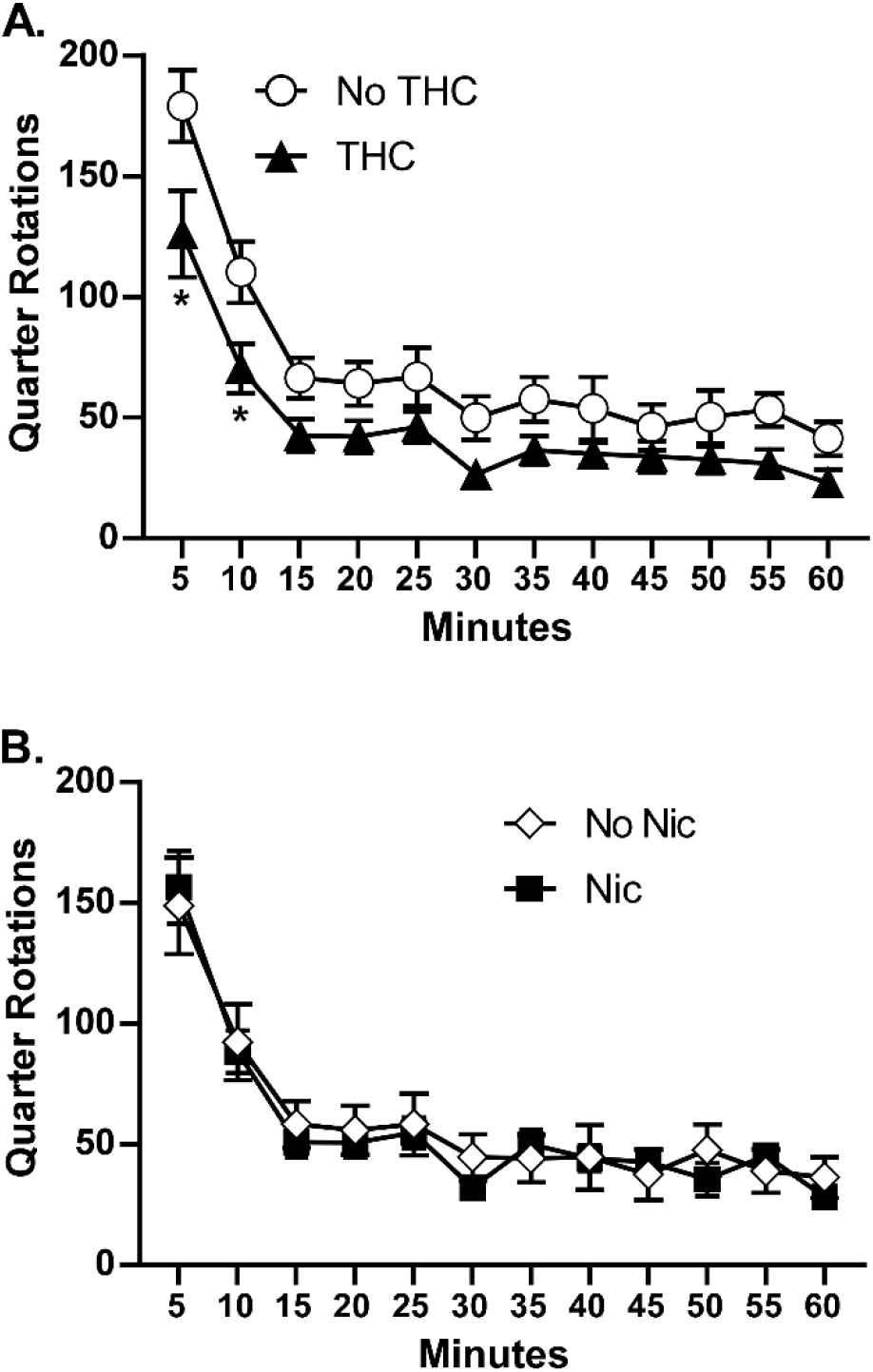
Baseline Wheel Activity PND 150-151. Mean (N=16; ±SEM) quarter rotations of the wheel for the groupings that received THC or no THC (A) or that received Nicotine or no Nicotine (B). A significant difference between groups is indicated with *.

### 3.4 Experiment 3: Effect of repeated adolescent nicotine and THC inhalation on volitional nicotine vapor exposure

The rats that were exposed to THC (alone or in combination with nicotine) obtained fewer vapor deliveries than those without THC exposure (PG and Nic alone) by the end of acquisition (**Figure 5**). The unavoidable interruption after Session 9 due to facilities emergency was followed by a return to approximately the same number of vapor deliveries in Sessions 10-12. The initial three factor ANOVA confirmed significant effects of Session [F (8, 224) = 2.86; P=0.005] and the interaction of Session with the presence/absence of THC [F (8, 224) = 1.98; P<0.05], of the interaction of Session with the presence/absence of Nicotine [F (8, 224) = 3.42; P<0.005], and the interaction of all three factors [F (8, 224) = 2.16; P<0.05] on vapor deliveries. The follow-up analysis of the impact of THC confirmed significant effects of Session [F (11, 330) = 3.26; P<0.0005] and the interaction of adolescent vapor condition with Session [F (11, 330) = 1.94; P<0.05], as did the analysis for the impact of nicotine [Session: F (11, 330) = 3.34; P<0.0005; interaction: F (11, 330) = 2.76; P<0.005]. The follow-up Dunnett post-hoc analysis of the impact of adolescent THC confirmed that compared with the first session, fewer deliveries were obtained by the No THC group on Sessions 2-8, 11-12 and by the THC group on Sessions 12. The follow-up Dunnett post-hoc analysis of the impact of adolescent nicotine confirmed that compared with the first session, fewer deliveries were obtained by the No Nicotine group on Sessions 3-6, 12 and by the Nicotine group on Sessions 8, 10-12.

**Figure 5:**
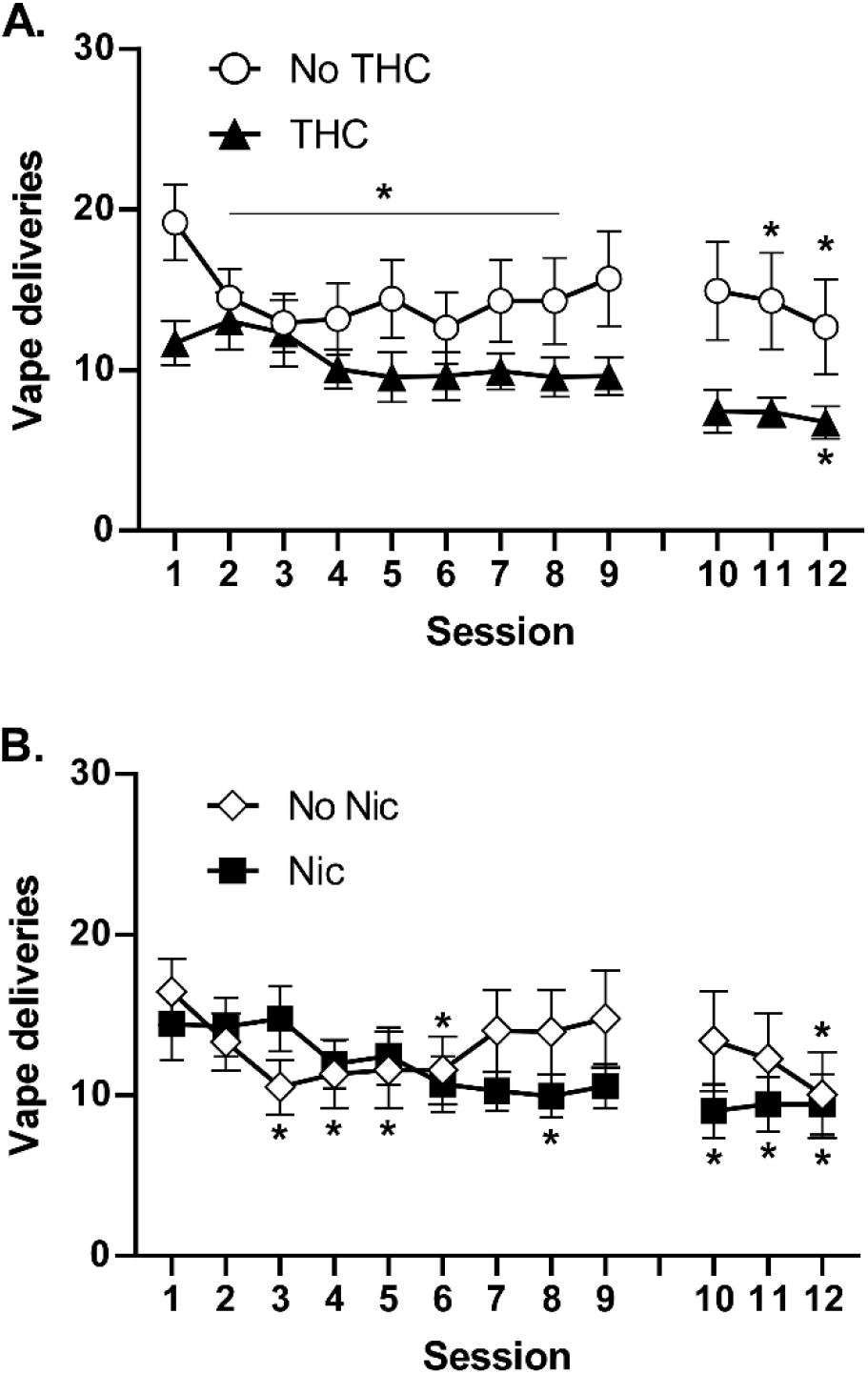
Acquisition of nicotine vapor (30 mg/mL) self-administration. Mean (N=16; ±SEM) Vapor Deliveries obtained during acquisition for the groupings that received THC or no THC (A) or that received Nicotine or no Nicotine (B). A significant difference from the first session within group is indicated with *. Animals were idled for three weeks between sessions 9 and 10 due to an unavoidable facilities interruption.

In the FR 5 experiment, the THC-exposed rats obtained fewer vapor deliveries (**Figure 6A**) and the three factor ANOVA confirmed a significant effect of THC/noTHC condition [F (1, 28) = 5.90; P<0.05], of the FR condition [F (6, 168) = 16.90; P<0.0001], and of the interaction of THC with FR condition [F (6, 168) = 4.02; P=0.001] on vapor deliveries.

**Figure 6:**
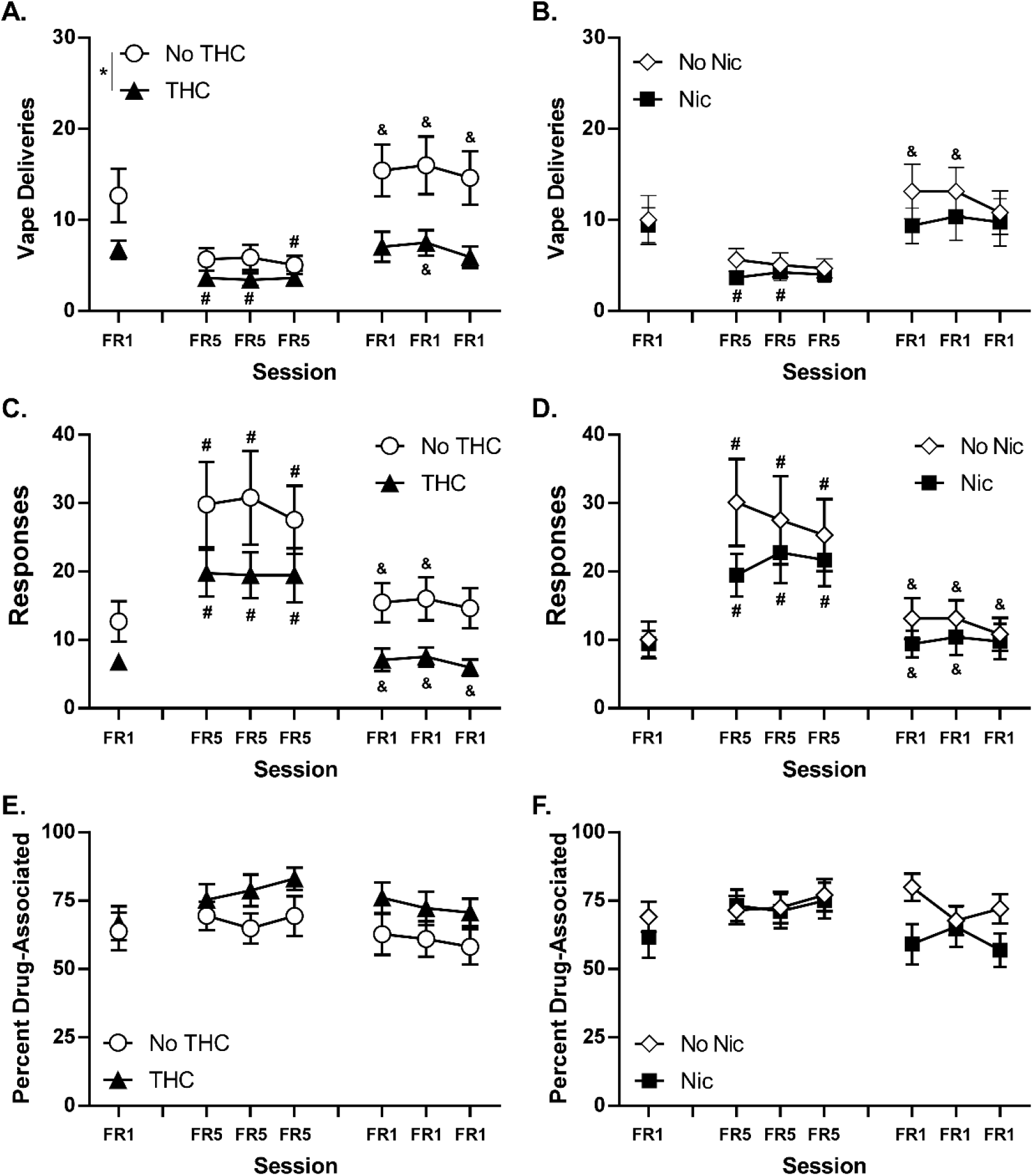
Nicotine vapor (30 mg/mL) self-administration. Mean (N=16; ±SEM) Vapor Deliveries (A, B), total Responses on the drug-associated manipulandum (C, D) and Percent of responses directed to the drug-associated manipulandum (E, F) for the groupings that received THC or no THC (A, C, E) or that received Nicotine or no Nicotine (B, D, F). A significant difference between groups is indicated with *. Within group, significant differences from the initial FR1 session are indicated with # and from the third FR5 session with &.

There were no significant effects of the adolescent nicotine factor alone or in interaction with other factors. Post-hoc analysis of the FR factor after the follow up two factor ANOVA, collapsed across all groups, confirmed that significantly fewer vapor deliveries were obtained in all three FR5 sessions compared with the FR1 baseline and compared with all three post-FR5 FR1 sessions.

In the follow-up analysis of the THC factor, the Tukey post-hoc analysis further confirmed that within the no-THC group fewer vape deliveries were obtained on the third FR5 session compared with the FR1 baseline. In addition, significantly fewer vape deliveries were obtained in all three FR5 sessions compared with all three of the following FR1 sessions. Intakes also differed significantly between the second and third FR5 sessions.

Within the THC group, significantly fewer vape deliveries were obtained in the first and second FR5 sessions compared with the FR1 baseline. In addition, significantly more vape deliveries were obtained in the second FR1 session after the FR5 sessions compared with all three FR5 sessions.

Analysis of the total drug-associated responses with a three factor ANOVA confirmed a significant effect of FR condition [F (6, 168) = 21.94; P<0.0001] but not of either adolescent treatment factor (**Figure 6C**). Post-hoc analysis of the FR factor after the follow up two factor ANOVA collapsed across all groups, confirmed that significantly more responses were emitted in all three FR5 sessions compared with the FR1 baseline and compared with all three post-FR5 FR1 sessions.

Analysis of the percent of responses directed at the drug-associated manipulandum (**Figure 6E**) with a three factor mixed-effects analysis confirmed a significant effect of THC condition [F (1, 28) = 5.16; P<0.05] and of the interaction of THC with Nicotine factor [F (1, 28) = 15.47; P<0.0005]; no significant effects of FR condition were confirmed. The Tukey post-hoc analysis comparing all four original groups confirmed that the Nicotine exposed animals exhibited a significantly lower percentage of responses on the drug-associated manipulandum compared with the PG and Combination groups.

Follow-up analysis of the adolescent nicotine versus no-nicotine groups did not confirm any significant effect of this factor on vape deliveries, total responses, or percent drug-associated responses.

### 3.4 Experiment 4: Effect of heroin injection on nociception and body temperature

Heroin injection significantly slowed tail-withdrawal latency in a dose- and time-dependent manner (**Figure 7**). The 3-factor analyses focused on the effect of time after injection for each dose. The analyses confirmed significant effects of time after vehicle [F (3, 56) = 10.89; P<0.0001] (not shown; range 2.1-5.1 30-60 minutes post-injection), 0.56 mg/kg [F (3, 56) = 52.61; P<0.0001], 1.0 mg/kg [F (3, 56) = 72.17; P<0.0001], and 1.56 mg/kg [F (3, 56) = 178.1; P<0.0001] heroin injection. There was a significant effect of the Nicotine/No nicotine factor for the 1.0 mg/kg dose [F (1, 56) =6.64; P<0.05] and an interaction of the THC/No THC factor with time [F (3, 56) = 3.20; P<0.05] after the 0.56 mg/kg heroin injection. Follow up post-hoc analysis of the THC/noTHC groupings for the 0.56 mg/kg dose confirmed a significant difference 30 minutes after injection and the follow-up post-hoc analysis of the Nicotine/no Nicotine groupings for the 1.0 mg/kg dose confirmed a significant difference 60 minutes after injection.

**Figure 7:**
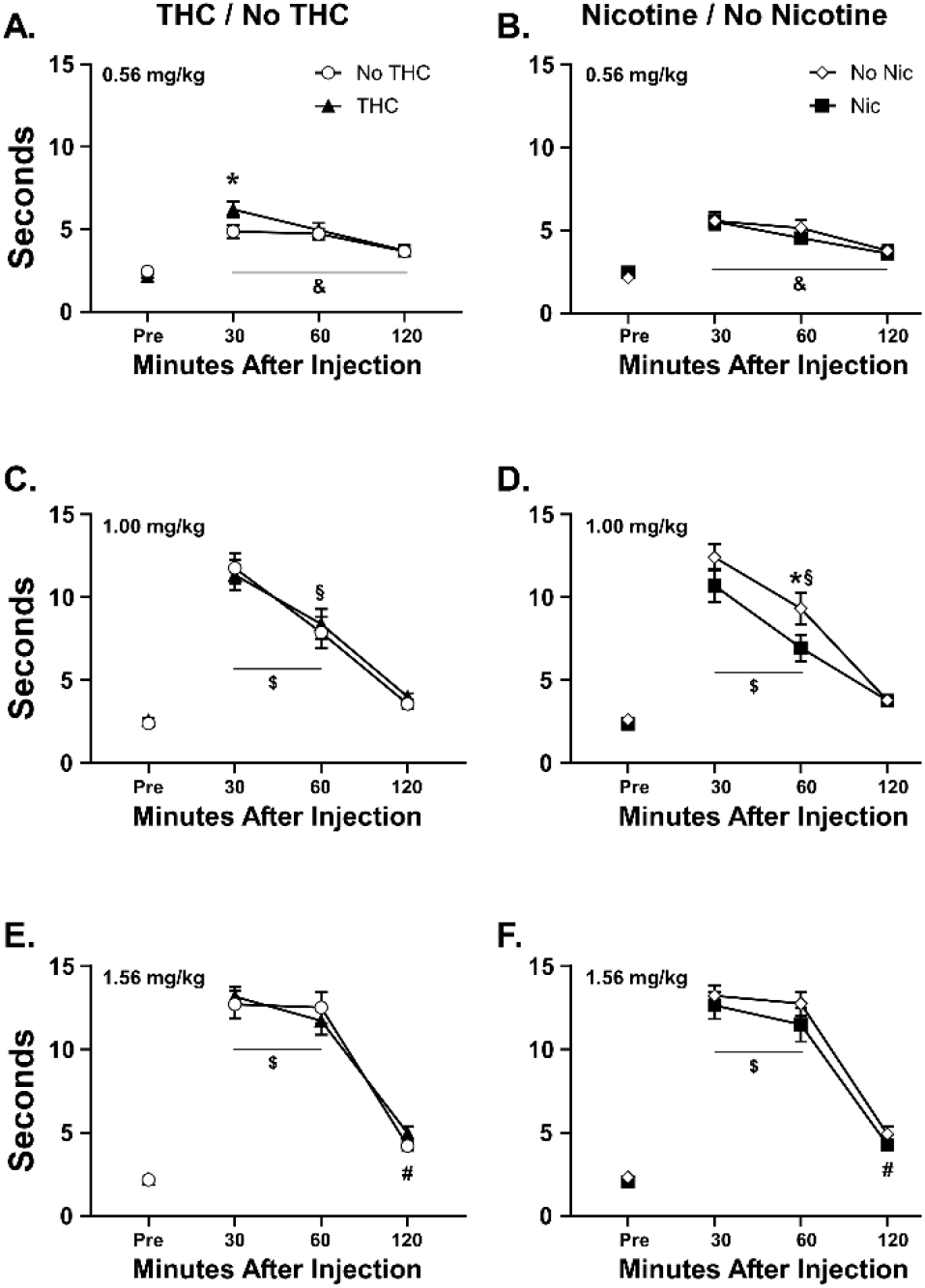
Mean (N=16 per group; ±SEM) tail-withdrawal latencies following injection with heroin (0.56-1.56 mg/kg, s.c.) are depicted for groups which did or did not receive THC (A., C., E.) and for groups which did or did not receive nicotine (B., D., F.). A significant difference between groups is indicated with *. Across group, a significant difference from all other time points is indicated with &. A significant difference from the baseline is indicated with #. A significant difference from the baseline and 120 minutes is indicated with $.

There were no significant effects of heroin injection on body temperature confirmed in the analysis (not shown).

### 3.5 Experiment 5: Effect of repeated adolescent nicotine and THC inhalation on volitional heroin vapor exposure

Responding for nicotine vapor under the FR1 response contingency after the break for the heroin anti-noiciception study approximated the same levels (**Figure 8**) as initially expressed at the end of the FR5/FR1 study (**Figure 6**). Vape deliveries were unaffected by the addition of menthol, a change to the 60 mg/mL concentration and a change of operant box for each animal. The group differences remained consistent with the adolescent THC exposure groups obtaining fewer vapor deliveries than the groups that did not receive adolescent THC. The analysis confirmed a significant effect of THC/No-THC group [F (1, 29) = 6.11; P<0.05] and of Session [F (5.027, 145.8) = 2.39; P<0.05], but the post-hoc test did not further confirm any specific differences. No differences were observed between the nicotine and no-nicotine adolescent treatment groups.

**Figure 8:**
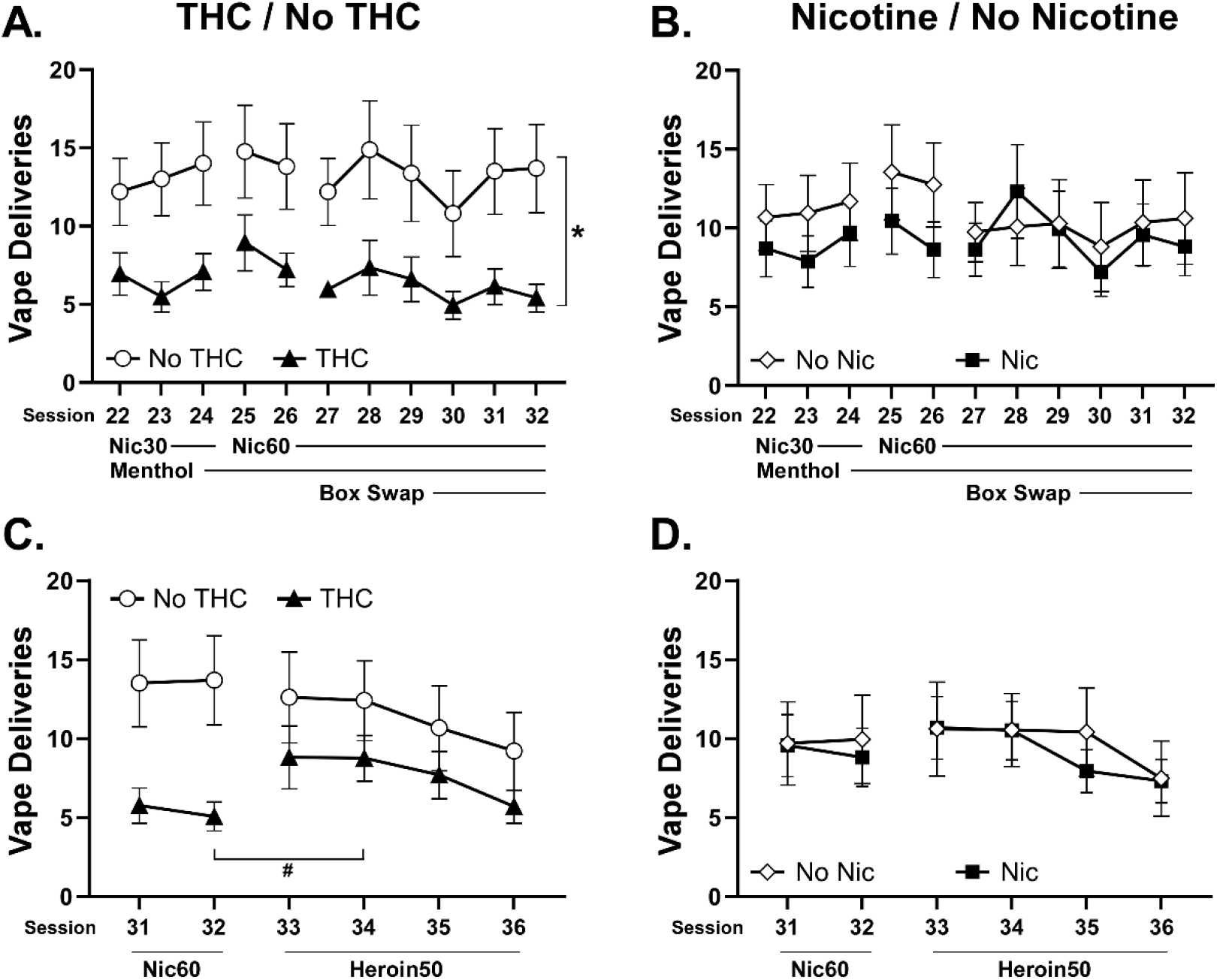
Mean (±SEM) vapor deliveries of Nicotine (30, 60 mg/mL) and Heroin (60 mg/mL) obtained by groups exposed as adolescents to PG, THC, Nicotine or THC+Nicotine vapor, grouped as the THC vs no-THC groups, and as the Nicotine versus no-Nicotine groups. Sessions 31 and 32 are repeated in the lower panels for assessment of the change when heroin was introduced. A difference between sessions within group is indicated by # and a difference between groups, across sessions, is indicated with *.

The introduction of heroin (50 mg/mL) vapor as the reinforcer significantly increased vapor deliveries obtained by the THC-exposed groups. The analysis confirmed a significant interaction between group and Session [F (5, 146) = 2.54; P<0.05] and the post-hoc Dunnett test confirmed that, relative to the final nicotine vapor session, the THC-exposed group obtained significantly more vapor deliveries on the second heroin session.

Nociception was assessed on the first day of heroin vapor availability. An anti-nociceptive effect was confirmed (Pre/Post: F (1, 27) = 79.45; P<0.0001) and the post-hoc test further confirmed that latencies were significantly slower after self-administration in all four adolescent treatment groups (**Figure 9**). No differences between groups were confirmed.

**Figure 9:**
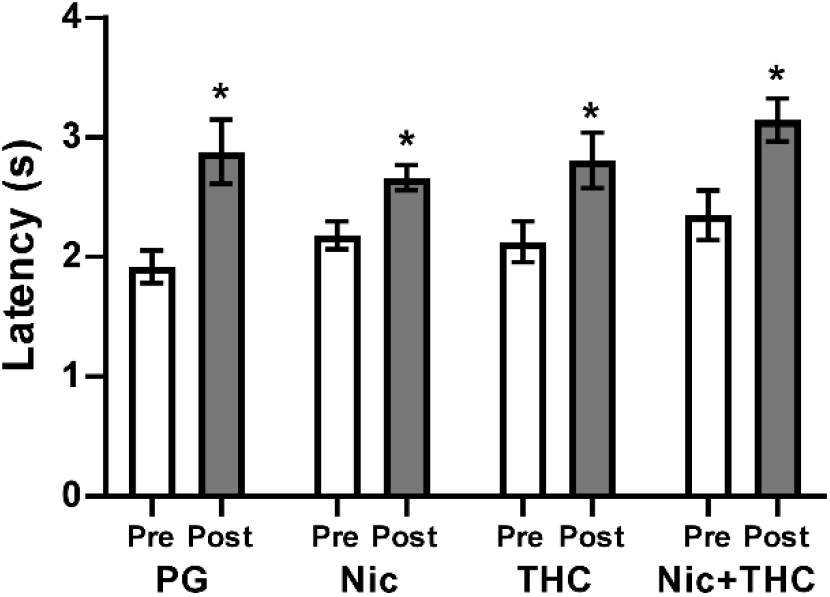
Mean (N=8 per group; ±SEM) tail-withdrawal latencies before and after the first heroin vapor self-administration session are depicted for adolescent treatment groups. A significant difference between the Pre- and Post-session measurements is indicated with *.

## Discussion

The study demonstrates that there are physiological and behavioral effects of repeated adolescent exposure to Δ^9^-tetrahydrocannabinol (THC) via Electronic Drug Delivery Systems (EDDS) vapor inhalation that last into adulthood in female rats. Vapor exposure to THC produced a lasting tolerance to the temperature disrupting effects of THC, reduced spontaneous activity on an exercise wheel and decreased volitional responding for nicotine vapor. In contrast, there were no lasting differences in the baseline wheel activity levels or in the effects of acute nicotine pre-treatment associated with the adolescent exposure to nicotine vapor. This is unlike our prior result for adolescent females exposed to twice daily nicotine vapor inhalation PND 31-40 using the 30 mg/mL concentration (Gutierrez et al., 2022), which suggests the potential importance of dose to the outcome. The addition of nicotine to the THC during adolescent exposure did not significantly modify the impact of the THC.

### Wheel Activity and Thermoregulation

The rats exposed to nicotine (alone or in combination with THC) had statistically indistinguishable wheel activity compared with the rats which did not receive adolescent nicotine, in the baseline test as well as in both of the PG and Nicotine inhalation conditions. In contrast, the rats that were exposed to THC (alone or in combination with nicotine) exhibited less baseline wheel activity as well as less activity under the PG challenge condition. The effect of nicotine, delivered by vapor inhalation, was to suppress wheel activity. This was similar to a suppressive effect of 0.4-0.8 mg/kg nicotine, s.c., injection on wheel activity previously reported (Bryson et al., 1981; Gutierrez et al., 2022). Twice daily adolescent vapor exposure to THC on 10 consecutive days (PND 36-45) produced lasting tolerance the thermoregulatoryeffects of THC in young adulthood, similar to a prior study with rats treated twice daily PND 35-39, 42-46 (Nguyen et al., 2020b). Similar thermoregulatory tolerance after twice daily repeated THC vapor exposure also has been reported for adult female rats (Nguyen et al., 2020a; Nguyen et al., 2018). This study extended those observations to show additional lasting behavioral consequences include reductions in spontaneous wheel activity.

As one minor caveat, one recent report suggest that vaping nicotine and THC together leads to lower plasma drug levels than when the same concentration of either drug is vaped alone (Breit et al., 2022). However given that the magnitude of tolerance to hypothermic effects of THC was similar across the THC-only and THC-Nicotine groups, and similar to our prior report (Nguyen et al., 2016b), this seems unlikely to have occurred in the present study.

### Volitional nicotine vapor exposure

The rats responded for nicotine in a manner consistent with some criteria for self-administration, as this term is used to describe intentional drug seeking behavior to an individually determined level of intoxication. Session intake under FR1 contingency was relatively consistent from Session 2 to Session 32 on a group mean basis. Continued drug-taking is a first-principle of self-administration. The declining trends sometimes associated with sequential days of intravenous nicotine self-administration (O’Dell and Koob, 2007) were not observed, presumably prevented with the intermittent schedule of sessions. Group means exceeded a 2:1 ratio of drug-associated : non-associated responses, a criterion which has been used to infer self-administration in some studies (Freels et al., 2020; Spencer et al., 2018; Stringfield et al., 2023). Total drug-associated responses increased significantly after the introduction of FR5 and declined again after the restoration of FR1. These are similar to effects previously reported for female rats in a prior study (Gutierrez et al., 2022) and are, together, highly consistent with drug-seeking behavior. This finding supports prior work with different vapor delivery and methodological approaches which also showed nicotine self-administration by vapor inhalation (Cooper, Akers and Henderson, 2021; Lallai et al., 2021; Smith et al., 2020).

The present study found that adolescent THC exposure *reduces* the responding for nicotine vapor. This appeared to be a preference set-point, since responding increased under the FR5 contingency and the percentage of drug-associated responses was, if anything, slightly higher than for that of the non-THC-exposed rats. Interestingly, the addition of menthol to the nicotine vapor did not affect drug seeking behavior (**Figure 8**), unlike a prior investigation in mice (Cooper, Akers and Henderson, 2021), nor did the increase in nicotine concentration. It is possible that the former was due to the nicotine doses being at the higher end of the dose range and therefore no additional effect of menthol could be detected. The lack of difference when the vapor concentration was increased to 60 mg/mL in the PG is perhaps unexpected but the dose-effect curves for intravenous self-administration of nicotine can be quite flat, e.g., across 2-to 3-fold differences in dose. For example, female mice self-administered similar intravenous infusions of nicotine whether it was 0.3 or 0.1 mg/kg/infusion (Dukes et al., 2020) and male rats self-administered similar numbers of intravenous infusions of 0.03 or 0.06 mg/kg/infusion nicotine (O’Dell et al., 2007). It is therefore most likely that the 30 mg/mL and 60 mg/mL conditions used here did not produce a large enough effective difference in delivered dose.

The adolescent nicotine inhalation did not affect adult responding for nicotine vapor, whereas adolescent THC exposure reduced nicotine self-administration in this study. This is similar to an effect reported in female mice injected daily during adolescence with nicotine, the CB1 full agonist WIN55,212-2 (WIN) or the combination (Dukes et al., 2020), in which adult intravenous nicotine self-administration was lower than the control group in the WIN or WIN+Nicotine groups, but unchanged in the nicotine-only group. Interestingly, male mice exposed as adolescents to WIN alone or in combination with nicotine self-administered more nicotine in adulthood in that study. In all of these cases the group effects were observed at the lowest unit doses of nicotine in a dose-substitution assessment, i.e., at or below the training dose, something that was not assessed in this study.

### Volitional heroin vapor exposure

The difference in volitional responding for nicotine vapor that was associated with adolescent THC exposure appeared to be selective for nicotine. The THC groups obtained more vapor deliveries when heroin was introduced, and this resulted in closing the persisting difference with the no-THC groups. The impact of the volitional heroin vapor exposure on the involuntary anti-nociceptive effect was similar (**Figure 9**), further emphasizing the similar level of subjective intoxication. This, combined with the differential effect of adolescent THC versus Nicotine vapor exposure on the anti-nociceptive effects of THC, underlines the selectivity of the insult, depending on the drug. Prior studies have shown repeated adolescent injections of THC lead to increased heroin self-administration in acquisition (Ellgren, Spano and Hurd, 2007; Lecca et al., 2020), or in yohimbine-induced re-instatement after no difference in acquisition (Stopponi et al., 2014), in adulthood. Adult female rats exposed as adolescents to THC vapor expressed higher rates of self-administration of fentanyl at low unit doses, after no difference in the acquisition of oxycodone self-administration (Nguyen et al., 2020b). Although the primary focus here was on nicotine self-administration, the heroin vapor experiment further supports the findings that repeated adolescent THC may lead to a lasting vulnerability to the reinforcing effects of heroin.

### Assessing Risks into Middle Age

It is notable that these studies successfully assessed drug seeking behavior into middle age in the rat, with the final heroin vapor self-administration studies completed around 50 weeks of age. Thus it is possible to use rat models to investigate age-related issues that may further understanding of a recent increase in middle aged opioid-related fatalities (Monnat, 2022). Controlled animal models are needed to parse factors that may explain differential rates in men versus women, or in ethnic and racial populations, by isolating the effects of drug exposure in the absence of human social factors. This latter is particularly critical given long term NIH funding disparities (Lauer and Roychowdhury, 2021; Taffe and Gilpin, 2021b) which leave topics of concern to Black PIs, across many neuropsychiatric domains (Gilpin and Taffe, 2021; Harnett, 2020; Lauer et al., 2021; Taffe, 2021) including substance use disorders (Acevedo et al., 2018; CPDDBoard, 2022; Taffe and Gilpin, 2021a), at a significant disadvantage (Hoppe et al., 2019). Here, we were able to isolate differential adolescent exposure and then determine lasting consequences throughout the young adult to middle age interval with all subsequent drug exposure similar across these groups. In longitudinal human studies, it would be far more likely that adolescent drug exposure is associated with additional significant differences in drug exposure throughout the early to middle adult age range, complicating inference about the adolescent drug exposure. It should be additionally noted on a practical level that the use of the vapor inhalation model for self-administration overcame typical subject loss due to, e.g., obstructed catheters and catheter-related health problems, that might be expected in an intravenous self-administration approach.

Overall, these data do not illustrate significant lasting consequences of repeated adolescent nicotine exposure by vapor inhalation on activity patterns on an exercise wheel, nor on nicotine or heroin vapor self-administration. This was the case when nicotine was administered alone or in the context of coincident THC vapor inhalation. The THC exposure, in contrast, produced lasting consequences as was the case in our prior studies (Nguyen et al., 2020b; Nguyen et al., 2018). In this case tolerance to the hypothermic effects of acute THC, lower activity on the running wheel, and reduced nicotine vapor self-administration were observed in adulthood; these effects were not significantly modulated by the addition of nicotine during repeated adolescent exposure.

## Acknowledgements

*These studies were supported by the Tobacco Related Disease Research Program (T31IP1832 and T33IR6653, MAT), a UCSD Chancellor’s Post-doctoral Fellowship (AG), UCSD IRACDA (AG) and the NIH (R01 DA042211, MAT). None of the funding bodies had any influence on the study design, data interpretation, manuscript creation or in the decision of when and what to publish from the studies conducted. Data will be made available to qualified individuals upon legitimate request*.

## Literature Cited

Acevedo, A., Panas, L., Garnick, D., Acevedo-Garcia, D., Miles, J., Ritter, G., Campbell, K., 2018. Disparities in the Treatment of Substance Use Disorders: Does Where You Live Matter? J Behav Health Serv Res 45(4), 533–549, doi: 10.1007/s11414-018-9586-y.

Blundell, M., Dargan, P., Wood, D., 2018. A cloud on the horizon-a survey into the use of electronic vaping devices for recreational drug and new psychoactive substance (NPS) administration. QJM 111(1), 9–14, doi: 10.1093/qjmed/hcx178.

Breit, K.R., Rodriguez, C.G., Hussain, S., Thomas, K.J., Zeigler, M., Gerasimidis, I., Thomas, J.D., 2022. A Model of Combined Exposure to Nicotine and Tetrahydrocannabinol via Electronic Cigarettes in Pregnant Rats. Front Neurosci 16, 866722, doi: 10.3389/fnins.2022.866722.

Bryson, R., Biner, P.M., McNair, E., Bergondy, M., Abrams, O.R., 1981. Effects of nicotine on two types of motor activity in rats. Psychopharmacology 73(2), 168–170, doi:

Cooper, S.Y., Akers, A.T., Henderson, B.J., 2021. Flavors Enhance Nicotine Vapor Self-administration in Male Mice. Nicotine & tobacco research : official journal of the Society for Research on Nicotine and Tobacco 23(3), 566–572, doi: 10.1093/ntr/ntaa165.

CPDDBoard, 2022. Funding Inequities Letter to NIH. https://cpdd.org/funding-inequities-letter-to-nih/.

Dukes, A.J., Fowler, J.P., Lallai, V., Pushkin, A.N., Fowler, C.D., 2020. Adolescent Cannabinoid and Nicotine Exposure Differentially Alters Adult Nicotine Self-Administration in Males and Females. Nicotine Tob Res 22(8), 1364–1373, doi: 10.1093/ntr/ntaa084.

Ellgren, M., Spano, S.M., Hurd, Y.L., 2007. Adolescent cannabis exposure alters opiate intake and opioid limbic neuronal populations in adult rats. Neuropsychopharmacology : official publication of the American College of Neuropsychopharmacology 32(3), 607–615, doi:

Fowler, C.D., Gipson, C.D., Kleykamp, B.A., Rupprecht, L.E., Harrell, P.T., Rees, V.W., Gould, T.J., Oliver, J., Bagdas, D., Damaj, M.I., Schmidt, H.D., Duncan, A., De Biasi, M., Basic Science Network of the Society for Research on, N., Tobacco, 2018. Basic Science and Public Policy: Informed Regulation for Nicotine and Tobacco Products. Nicotine & tobacco research : official journal of the Society for Research on Nicotine and Tobacco 20(7), 789–799, doi: 10.1093/ntr/ntx175.

Freels, T.G., Baxter-Potter, L.N., Lugo, J.M., Glodosky, N.C., Wright, H.R., Baglot, S.L., Petrie, G.N., Yu, Z., Clowers, B.H., Cuttler, C., Fuchs, R.A., Hill, M.N., McLaughlin, R.J., 2020. Vaporized Cannabis Extracts Have Reinforcing Properties and Support Conditioned Drug-Seeking Behavior in Rats. J Neurosci 40(9), 1897–1908, doi: 10.1523/JNEUROSCI.2416-19.2020.

Garber, J.C., Barbee, R.W., Bielitzki, J.T., Clayton, L.A., Donovan, J.C., Hendriksen, C.F.M., Kohn, D.F., Lipman, N.S., Locke, P.A., Melcher, J., Quimby, F.W., Turner, P.V., Wood, G.A., Wurbel, H., 2011. Guide for the Care and Use of Laboratory Animals, 8th Edition. National Academies Press, Washington D.C.

Gilpin, N.W., Taffe, M.A., 2021. Toward an Anti-Racist Approach to Biomedical and Neuroscience Research. J Neurosci 41(42), 8669–8672, doi: 10.1523/JNEUROSCI.1319-21.2021.

Gilpin, N.W., Wright, M.J., Jr., Dickinson, G., Vandewater, S.A., Price, J.U., Taffe, M.A., 2011. Influences of activity wheel access on the body temperature response to MDMA and methamphetamine. Pharmacology, biochemistry, and behavior 99(3), 295–300, doi: 10.1016/j.pbb.2011.05.006.

Gutierrez, A., Creehan, K.M., Taffe, M.A., 2021. A vapor exposure method for delivering heroin alters nociception, body temperature and spontaneous activity in female and male rats. J Neurosci Methods 348, 108993, doi: 10.1016/j.jneumeth.2020.108993.

Gutierrez, A., Nguyen, J.D., Creehan, K.M., Grant, Y., Taffe, M.A., 2022. Adult consequences of repeated nicotine vapor inhalation in adolescent rats. bioRxiv, 2022.2011.2017.516984, doi: 10.1101/2022.11.17.516984.

Harnett, N.G., 2020. Neurobiological consequences of racial disparities and environmental risks: a critical gap in understanding psychiatric disorders. Neuropsychopharmacology : official publication of the American College of Neuropsychopharmacology 45(8), 1247–1250, doi: 10.1038/s41386-020-0681-4.

Henderson, B.J., Cooper, S.Y., 2021. Nicotine formulations impact reinforcement-related behaviors in a mouse model of vapor self-administration. Drug and alcohol dependence 224, 108732, doi: 10.1016/j.drugalcdep.2021.108732.

Hoppe, T.A., Litovitz, A., Willis, K.A., Meseroll, R.A., Perkins, M.J., Hutchins, B.I., Davis, A.F., Lauer, M.S., Valantine, H.A., Anderson, J.M., Santangelo, G.M., 2019. Topic choice contributes to the lower rate of NIH awards to African-American/black scientists. Sci Adv 5(10), eaaw7238, doi: 10.1126/sciadv.aaw7238.

Hussain, S., Breit, K.R., Thomas, J.D., 2022. The effects of prenatal nicotine and THC E-cigarette exposure on motor development in rats. Psychopharmacology 239(5), 1579–1591, doi: 10.1007/s00213-022-06095-8.

Javadi-Paydar, M., Creehan, K.M., Kerr, T.M., Taffe, M.A., 2019a. Vapor inhalation of cannabidiol (CBD) in rats. Pharmacology, biochemistry, and behavior 184, 172741, doi: 10.1016/j.pbb.2019.172741.

Javadi-Paydar, M., Kerr, T.M., Harvey, E.L., Cole, M., Taffe, M.A., 2019b. Effects of nicotine and THC vapor inhalation administered by an electronic nicotine delivery system (ENDS) in male rats. Drug and alcohol dependence 198, 54–62, doi: 10.1016/j.drugalcdep.2019.01.027.

Johnston, L.D., Miech, R.A., O’Malley, P.M., Bachman, J.G., Schulenberg, J.E., Patrick, M.E., 2021. Monitoring the Future national survey results on drug use, 1975-2020: 2020 Overview Key Findings on Adolescent Drug Use. http://www.monitoringthefuture.org/pubs/monographs/mtf-overview2020.pdf.

Lallai, V., Chen, Y.C., Roybal, M.M., Kotha, E.R., Fowler, J.P., Staben, A., Cortez, A., Fowler, C.D., 2021. Nicotine e-cigarette vapor inhalation and self-administration in a rodent model: Sex- and nicotine delivery-specific effects on metabolism and behavior. Addict Biol 26(6), e13024, doi: 10.1111/adb.13024.

Lauer, M.S., Doyle, J., Wang, J., Roychowdhury, D., 2021. Associations of topic-specific peer review outcomes and institute and center award rates with funding disparities at the National Institutes of Health. Elife 10, doi: 10.7554/eLife.67173.

Lauer, M.S., Roychowdhury, D., 2021. Inequalities in the distribution of National Institutes of Health research project grant funding. Elife 10, doi: 10.7554/eLife.71712.

Lecca, D., Scifo, A., Pisanu, A., Valentini, V., Piras, G., Sil, A., Cadoni, C., Di Chiara, G., 2020. Adolescent cannabis exposure increases heroin reinforcement in rats genetically vulnerable to addiction. Neuropharmacology 166, 107974, doi: 10.1016/j.neuropharm.2020.107974.

Miech, R.A., Johnston, L.D., O’Malley, P.M., Bachman, J.G., Schulenberg, J.E., Patrick, M.E., 2022. Monitoring the Future national survey results on drug use, 1975-2021. Volume I, Secondary school students

Miller, M.L., Moreno, A.Y., Aarde, S.M., Creehan, K.M., Vandewater, S.A., Vaillancourt, B.D., Wright, M.J., Jr., Janda, K.D., Taffe, M.A., 2013. A methamphetamine vaccine attenuates methamphetamine-induced disruptions in thermoregulation and activity in rats. Biol Psychiatry 73(8), 721–728, doi: 10.1016/j.biopsych.2012.09.010.

Monnat, S.M., 2022. Demographic and Geographic Variation in Fatal Drug Overdoses in the United States, 1999-2020. Ann Am Acad Pol Soc Sci 703(1), 50–78, doi: 10.1177/00027162231154348.

Montanari, C., Kelley, L.K., Kerr, T.M., Cole, M., Gilpin, N.W., 2020. Nicotine e-cigarette vapor inhalation effects on nicotine & cotinine plasma levels and somatic withdrawal signs in adult male Wistar rats. Psychopharmacology 237(3), 613–625, doi: 10.1007/s00213-019-05400-2.

Morris, J.D., Pebley, K., Little, M.A., 2023. Vaping Opioids: Should We Be Worried? Am J Health Promot, 8901171231193785, doi: 10.1177/08901171231193785.

Nguyen, J.D., Aarde, S.M., Cole, M., Vandewater, S.A., Grant, Y., Taffe, M.A., 2016a. Locomotor Stimulant and Rewarding Effects of Inhaling Methamphetamine, MDPV, and Mephedrone via Electronic Cigarette-Type Technology. Neuropsychopharmacology : official publication of the American College of Neuropsychopharmacology 41(11), 2759–2771, doi: 10.1038/npp.2016.88.

Nguyen, J.D., Aarde, S.M., Vandewater, S.A., Grant, Y., Stouffer, D.G., Parsons, L.H., Cole, M., Taffe, M.A., 2016b. Inhaled delivery of Delta(9)-tetrahydrocannabinol (THC) to rats by e-cigarette vapor technology. Neuropharmacology 109, 112–120, doi: 10.1016/j.neuropharm.2016.05.021.

Nguyen, J.D., Creehan, K.M., Grant, Y., Vandewater, S.A., Kerr, T.M., Taffe, M.A., 2020a. Explication of CB(1) receptor contributions to the hypothermic effects of Delta(9)-tetrahydrocannabinol (THC) when delivered by vapor inhalation or parenteral injection in rats. Drug and alcohol dependence 214, 108166, doi: 10.1016/j.drugalcdep.2020.108166.

Nguyen, J.D., Creehan, K.M., Kerr, T.M., Taffe, M.A., 2020b. Lasting effects of repeated (9) - tetrahydrocannabinol vapour inhalation during adolescence in male and female rats. British journal of pharmacology 177(1), 188–203, doi: 10.1111/bph.14856.

Nguyen, J.D., Grant, Y., Creehan, K.M., Hwang, C.S., Vandewater, S.A., Janda, K.D., Cole, M., Taffe, M.A., 2019. Delta(9)-tetrahydrocannabinol attenuates oxycodone self-administration under extended access conditions. Neuropharmacology 151, 127–135, doi: 10.1016/j.neuropharm.2019.04.010.

Nguyen, J.D., Grant, Y., Kerr, T.M., Gutierrez, A., Cole, M., Taffe, M.A., 2018. Tolerance to hypothermic and antinoceptive effects of 9-tetrahydrocannabinol (THC) vapor inhalation in rats. Pharmacology, biochemistry, and behavior 172, 33–38, doi: 10.1016/j.pbb.2018.07.007.

O’Dell, L.E., Chen, S.A., Smith, R.T., Specio, S.E., Balster, R.L., Paterson, N.E., Markou, A., Zorrilla, E.P., Koob, G.F., 2007. Extended access to nicotine self-administration leads to dependence: Circadian measures, withdrawal measures, and extinction behavior in rats. The Journal of pharmacology and experimental therapeutics 320(1), 180–193, doi: jpet.106.105270[pii]10.1124/jpet.106.105270.

O’Dell, L.E., Koob, G.F., 2007. ‘Nicotine deprivation effect’ in rats with intermittent 23-hour access to intravenous nicotine self-administration. Pharmacology, biochemistry, and behavior 86(2), 346–353, doi: S0091-3057(07)00020-2 [pii] 10.1016/j.pbb.2007.01.004.

Pelham, W.E., 3rd, Tapert, S.F., Gonzalez, M.R., McCabe, C.J., Lisdahl, K.M., Alzueta, E., Baker, F.C., Breslin, F.J., Dick, A.S., Dowling, G.J., Guillaume, M., Hoffman, E.A., Marshall, A.T., McCandliss, B.D., Sheth, C.S., Sowell, E.R., Thompson, W.K., Van Rinsveld, A.M., Wade, N.E., Brown, S.A., 2021. Early Adolescent Substance Use Before and During the COVID-19 Pandemic: A Longitudinal Survey in the ABCD Study Cohort. J Adolesc Health 69(3), 390–397, doi: 10.1016/j.jadohealth.2021.06.015.

Smith, L.C., Kallupi, M., Tieu, L., Shankar, K., Jaquish, A., Barr, J., Su, Y., Velarde, N., Sedighim, S., Carrette, L.L.G., Klodnicki, M., Sun, X., de Guglielmo, G., George, O., 2020. Validation of a nicotine vapor self-administration model in rats with relevance to electronic cigarette use. Neuropsychopharmacology : official publication of the American College of Neuropsychopharmacology 45(11), 1909–1919, doi: 10.1038/s41386-020-0734-8.

Spencer, S., Neuhofer, D., Chioma, V.C., Garcia-Keller, C., Schwartz, D.J., Allen, N., Scofield, M.D., Ortiz-Ithier, T., Kalivas, P.W., 2018. A Model of Delta(9)-Tetrahydrocannabinol Self-administration and Reinstatement That Alters Synaptic Plasticity in Nucleus Accumbens. Biol Psychiatry 84(8), 601–610, doi: 10.1016/j.biopsych.2018.04.016.

Stopponi, S., Soverchia, L., Ubaldi, M., Cippitelli, A., Serpelloni, G., Ciccocioppo, R., 2014. Chronic THC during adolescence increases the vulnerability to stress-induced relapse to heroin seeking in adult rats. Eur Neuropsychopharmacol 24(7), 1037–1045, doi: 10.1016/j.euroneuro.2013.12.012.

Stringfield, S.J., Sanders, B.E., Suppo, J.A., Sved, A.F., Torregrossa, M.M., 2023. Nicotine Enhances Intravenous Self-administration of Cannabinoids in Adult Rats. Nicotine & tobacco research : official journal of the Society for Research on Nicotine and Tobacco 25(5), 1022–1029, doi: 10.1093/ntr/ntac267.

Taffe, M.A., 2021. NIH research funding disparities affect diversity, equity and inclusion goals of the ACNP. Neuropsychopharmacology : official publication of the American College of Neuropsychopharmacology 46(5), 880–881, doi: 10.1038/s41386-021-00969-9.

Taffe, M.A., Creehan, K.M., Vandewater, S.A., Kerr, T.M., Cole, M., 2021a. Effects of Delta(9)-tetrahydrocannabinol (THC) vapor inhalation in Sprague-Dawley and Wistar rats. Experimental and clinical psychopharmacology 29(1), 1–13, doi: 10.1037/pha0000373.

Taffe, M.A., Gilpin, N.W., 2021a. The Funding is the Science: Racial Inequity of NIH Funding for Substance Use Disorder Topics Should Be Abolished. Drug and alcohol dependence 229, 109163, doi: 10.1016/j.drugalcdep.2021.109163.

Taffe, M.A., Gilpin, N.W., 2021b. Racial inequity in grant funding from the US National Institutes of Health. Elife 10, doi: 10.7554/eLife.65697.

Taffe, M.A., Nguyen, J.D., Vandewater, S.A., Grant, Y., Dickerson, T.J., 2021b. Effects of alpha-pyrrolidino-phenone cathinone stimulants on locomotor behavior in female rats. Drug and alcohol dependence 227, 108910, doi: 10.1016/j.drugalcdep.2021.108910.

